# Father Absence and Accelerated Reproductive Development

**DOI:** 10.1101/123711

**Authors:** Lauren Gaydosh, Daniel W. Belsky, Benjamin W. Domingue, Jason D. Boardman, Kathleen Mullan Harris

## Abstract

Evidence shows that girls who experience father absence in childhood experience accelerated reproductive development in comparison to peers with present fathers. One hypothesis advanced to explain this empirical pattern is genetic confounding, wherein gene-environment correlation (rGE) causes a spurious relationship between father absence and reproductive timing. We test this hypothesis by constructing polygenic scores for age at menarche and first birth using recently available genome wide association study results and molecular genetic data on a sample of non-Hispanic white females from the National Longitudinal Study of Adolescent to Adult Health. Young women’s accelerated menarche polygenic scores were unrelated to their exposure to father absence. In contrast, earlier first-birth polygenic scores tended to be higher in young women raised in homes with absent fathers. Nevertheless, father absence and the polygenic scores independently and additively predict reproductive timing. We find limited evidence in support of the gene-environment correlation hypothesis.

## Introduction

Early pubertal timing is associated with risk for adverse behavioral and health outcomes across the life course (Karapanou and Papadimitriou 2010; Mendle et al. 2009; Tamakoshi et al. 2011). Pubertal timing is influenced by both genetic and environmental factors (Ellis 2004; Ellis et al. 2011; Polderman et al. 2015). One environmental factor consistently associated with earlier pubertal timing is exposure to childhood adversity (J. Belsky et al. 1991; Ellis 2004), in particular, absence of the biological father from the home (Webster et al. 2014). There are two hypotheses commonly advanced to explain the association between absence of the biological father and early puberty. One hypothesis, derived from evolutionary theory (Trivers 1972), is that father absence triggers accelerated maturation in the developing organism (J. Belsky et al. 1991). An alternative hypothesis is that the association is an artifact of gene-environment correlation (Rowe 2002).

We report a molecular genetic test of this rGE hypothesis. Using results from a recent genome-wide association study of age at menarche (Elks et al. 2010), we quantify genetic influences on pubertal timing in a sample of female adolescents from the National Longitudinal Study of Adolescent to Adult Health (Add Health). We test if the genetics discovered in GWAS of menarcheal timing (a) predict timing of puberty in Add Health girls; (b) correlate with girls’ exposure to father absence; and (c) explain any association between father absence and early puberty. We also consider genetics discovered in GWAS of age at first birth (Barban et al. 2016) and repeat our analysis in Add Health girls who report non-white race/ethnicity.

## Background

### Early Puberty and Life Course Development

Pubertal timing is of central interest to demographers, as it signals the start of reproductive capacity. For girls, age at menarche is associated with a cascade of behaviors and events in the transition to adulthood that relate to fertility and health outcomes. Earlier menarche is associated with earlier sexual debut, first pregnancy and birth, and marriage (Kiernan 1977; Sandler et al. 1984; Udry 2008). These associations replicate across cultural contexts (Udry and Cliquet 1982). Earlier pubertal timing is also associated with greater risk taking behavior more generally (Igra and Irwin Jr. 1996; Patton and McMorris 2004). Age at menarche is negatively correlated with poor health outcomes including type 2 diabetes (He et al. 2010) cardiovascular disease risk (Feng et al. 2008; Lakshman et al. 2009; Remsberg et al. 2005), breast cancer (Stoll et al. 1994), and mortality (Charalampopoulos et al. 2014; Tamakoshi et al. 2011).

### Father Absence and Early Puberty

Rooted in life course and evolutionary frameworks, a robust literature has documented associations between early childhood environment and reproductive profiles across adolescence and into adulthood (see Ellis 2004 for a thorough review). In particular, children exposed to psychosocial stressors during sensitive periods of childhood exhibit accelerated reproductive trajectories, such as early puberty, early sexual debut, and early childbearing (J. Belsky et al. 1991; Browning et al. 2004; Foster et al. 2008; Graber et al. 1995; Wu and Martinson 1993).

One specific childhood stressor that has received considerable attention is father absence. Girls who grow up in households from which their biological father is absent are consistently observed to experience menarche at younger ages as compared to peers with present fathers (Alvergne et al. 2008; Bogaert 2008; Culpin et al. 2014; Hoier 2003; Quinlan 2003). Father absence is also associated with subsequent reproductive timing; girls with absent fathers have younger age of first sex, pregnancy, and birth (Anderson 2015; Ellis et al. 2003; Kiernan and Hobcraft 1997; Mendle et al. 2009; Moffitt et al. 1992).

Research in anthropology and human development hypothesizes that associations between early-life stressors, including father absence, and accelerated reproductive development reflect a causal process (J. Belsky et al. 1991; Chisholm et al. 1993; Draper and Harpending 1982; Ellis and Garber 2000; Ellis 2004; Ellis et al. 1999). In this causal process, father absence, either specifically or as one of a several features of early adversity, signals to the developing child that resources may be scarce or unpredictable. This signal, in turn, elicits a biological response in the form of increased allocation of current resources toward achieving reproductive maturity as soon as possible (J. Belsky et al. 1991; Chisholm et al. 1993; Ellis 2004). Consistent with this hypothesis, father absence, as well as other measures of early adversity, are associated with accelerated pubertal development (Foster et al. 2008; Mendle et al. 2009, 2015). We focus here on father absence because it is central to hypotheses of environmental causation, and because it is among the best-replicated environmental correlates of accelerated pubertal development (Webster et al. 2014).

### Gene-environment Correlation as an Explanation for Association between Father Absence and Early Puberty

In contrast to anthropological and human development hypotheses of environmental causation, behavioral genetics researchers have proposed a genetic hypothesis for why girls who grow up in father absent households tend to experience accelerated reproductive development. This genetic hypothesis is built on three sets of research findings. First, reproductive timing is influenced by genetic factors (Barban et al. 2016; Burt et al. 2006; B. Campbell and Udry 1995; Elks et al. 2010; He et al. 2009; Tither and Ellis 2008; Towne et al. 2005). A recent meta-analysis estimates heritability for reproductive traits at 0.32 (Polderman et al. 2015), and a study of age at menarche specifically estimates heritability at 0.49 (Towne et al. 2005). In other words, roughly one-half of the variation in age at menarche has been attributed to genetic variation. Second, girls who mature earlier also experience earlier sexual debut (Moore et al. 2014), union formation (Kiernan 1977), and childbearing (Rowe 2002; Sandler et al. 1984; Udry 2008). Third, unions formed at earlier ages and with early childbearing are more likely to be unstable (Booth and Edwards 1985; Bumpass and Sweet 1972). The genetic hypothesis derived from this evidence posits that genetic factors that accelerate pubertal timing place young women at risk for early childbearing within unstable unions, which in turn results in their daughters inheriting both genetics of early puberty and a father-absent environment (Hardy et al. 1998; Mendle et al. 2006; Rowe 2000, 2002). This genetic hypothesis conceptualizes the association between father absence and daughters’ early puberty as spurious, confounded by a “gene-environment correlation” (rGE). Related work suggests that the genetic confounding may operate by fathers passing on genes to their children that both influence their own reproductive and mating behavior, as well as their children’s pubertal development (Comings et al. 2002).

To date, empirical tests of genetic confounding are limited to family-based genetic studies. These studies compare phenotypic similarities between pairs of relatives with different degrees of genetic relatedness, such as monozygotic and dizygotic twins. They use these comparisons to partition phenotypic variance into genetic and environmental components, and to further isolate environmental variance that is “shared” between siblings in a family (Plomin et al. 2013). Behavioral genetic studies find little evidence for “shared” environmental influence on pubertal timing, and generally interpret this as evidence against environmental causation hypotheses because father absence is an environmental exposure shared by siblings (Mendle et al. 2006; Rowe 2000; Ryan 2015; Tither and Ellis 2008). The advent of genome-wide association studies and the discovery of molecular genetic influences on reproductive development now afford an opportunity to put the genetic hypothesis to a molecular test.

### Genome-wide Association Studies of Human Reproductive Development

Advances in genome science are now yielding molecular detail about genetic influences on human traits and behaviors. Genome-wide association studies (GWAS) are large-scale data-mining expeditions in the human genome. They correlate millions of genetic variants, called “single-nucleotide polymorphisms” (SNPs) with phenotypes in samples of tens or even hundreds of thousands of individuals. GWAS conducted of age at menarche (Day et al. 2015; Perry et al. 2014) and age at first birth (Barban et al. 2016) provide an opportunity to conduct a molecular test of the rGE hypothesis advanced to explain associations between father absence and timing of puberty. GWAS results can be used to parameterize predictive algorithms, called “polygenic scores”, which can be applied to genomic data from independent samples (D. W. Belsky and Israel 2014; Dudbridge et al. 2013). Using this approach, we conduct the first molecular test of the hypothesis of gene-environment correlation between genetic influences on reproductive development and father absence and we evaluate genetic confounding of associations between father absence and early puberty.

## Data and Methods

### Data

The National Longitudinal Study of Adolescent to Adult Health (Add Health) is an ongoing, nationally-representative longitudinal study of the social, behavioral, and biological linkages in health and developmental trajectories from early adolescence into adulthood. The cohort was drawn from a probability sample of 132 middle and high schools and is representative of American adolescents in grades 7-12 in 1994-1995. Since the start of the project, participants have been interviewed in home at four data collection waves (numbered I-IV), most recently in 2008. Data include measures of family structure, reproductive development, and genome-wide molecular genetic data (Harris 2010; Harris et al. 2013).

#### Reproductive Timing

Age at menarche was recorded from interviews conducted at Waves I-III. We use data from the interview wave at which a woman first reported having had her first menstruation. Age at menarche is measured in completed years. Average age at menarche in the analytic sample of unrelated non-Hispanic white women is 12.3 years. All women in the analytic sample completed menarche by age 20 years.

Age at first sex was recorded from interviews conducted across Waves I-IV. We use data from the interview wave at which a woman first reported having had vaginal intercourse. Age at first sex was recorded in month units at Add Health Waves I and II and in year units at Waves III and IV. We exclude from analysis data from 13 women who reported an age at first sex less than ten years. 4% of the sample did not report having sex during follow-up. For the remainder of unrelated non-Hispanic white women, mean age-at-first-sex is 17.3 years.

Age at first birth was recorded from interviews conducted across Waves I-IV. We use data from the interview wave at which a woman first reported having had her first live birth in the pregnancy/birth-history portion of the interview. Age at first birth is measured in months. We exclude from analysis data from two women who reported first birth before age 10 years. 64% of women in the analytic sample reported a live birth by Wave IV. Of this group, mean age at first birth is 26.2 years.

Age at menarche, first sex, and first birth are positively correlated (Table 1). Women who develop later in adolescence tend to have a later age at sexual debut, followed by a later age at first birth. Age at first sex and age at first birth are most strongly correlated (.38). In Figure 1, we present Kaplan-Meier estimates for the cumulative proportion of women experiencing each reproductive event by age. The mean age at which women experience a given event is calculated as the area under the Kaplan-Meier survival function. All non-Hispanic white women experience menarche by age 20, with a mean age of 12.3. 96% experience sexual debut by age 32, with a mean age for those who have sex of 17.3. Finally, 64% experience a first birth by age 32, and the mean age for those who have a first birth is 26.2.

**Table 1.** Correlation Matrix, Unrelated White Females, n=2,631.

|  | Mean(SD)/% | Age at menarche | Age at first sex | Age at first birth | Accelerated menarche PGS | Earlier first-birth PGS | Father absent |
| --- | --- | --- | --- | --- | --- | --- | --- |
| Age at menarche | 12.2 (1.3) |  |  |  |  |  |  |
| Age at first sex | 17.1 (3.5) | 0.12 |  |  |  |  |  |
| Age at first birth | 24.7 (4.1) | 0.11 | 0.36 |  |  |  |  |
| Accelerated menarche PGS | 0 (1) | -0.23 | -0.04 | -0.04 |  |  |  |
| Earlier first-birth PGS | 0 (1) | -0.09 | -0.18 | -0.19 | 0.13 |  |  |
| Father absent | 29% | -0.07 | -0.15 | -0.14 | 0.01 | 0.08 |  |

**Figure 1.**
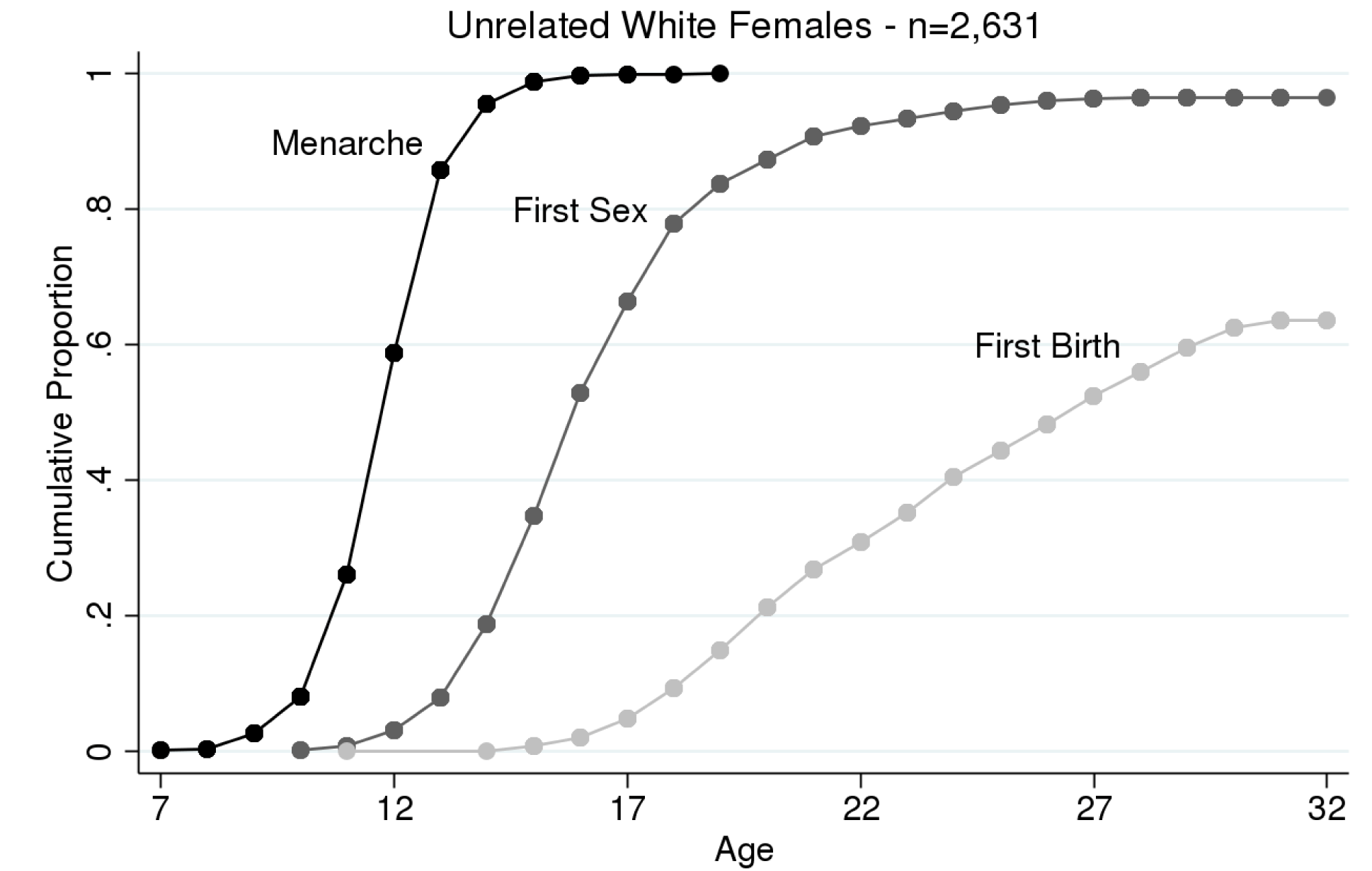
Reproductive Timing. Mean time to event: menarche = 12.3; first sex = 17.3; first birth = 26.

#### Father Absence

We measure father absence as an indicator for whether the child’s biological father was ever non-resident from birth to age seven. We construct this measure drawing from four sources of data: parent’s self-reported union history and current union status; youth’s reports of their current family structure; youth’s reports of the duration of life spent with current household members; and whether and for how long a youth lived with a nonresident biological parent (authors 2017). 18% of girls were born into a family structure where the biological father was non-resident. By age seven, 29% of girls experience biological father non-residence for a period of at least one year.

#### Genotyping

At the Wave IV interview in 2008, saliva and capillary whole blood were collected from respondents. 15,159 of 15,701 individuals interviewed consented to genotyping, and 12,254 agreed to genetic data archiving. After quality control procedures (Highland et al. 2017; McQueen et al. 2015), genotype data were available for 10,577 individuals. Data were generated for 8,266 individuals from the Illumina HumanOmni1-Quad chip and for 2,311 individuals from the Illumina HumanOmni2.5-Quad chip. We conducted analysis on the 631,990 single-nucleotide polymorphisms (SNPs) genotyped on both chips.

#### Polygenic Scores (PGS)

We calculate polygenic scores for Add Health participants based on published genome wide association study (GWAS) results for age at menarche from the ReproGen Consortium (Elks et al. 2010) and for age at first birth from the Social Science Genetic Association Consortium and Sociogenome (Barban et al. 2016). Scores were calculated following the method described by Dudbridge and colleagues (2013). Briefly, SNPs in the genotyped sample were matched to published GWAS results. For each of these SNPs (N=552,064 menarche, N=557,761 age at first birth), a loading was calculated as the number of phenotype-associated alleles multiplied by the effect-size estimated in the original GWAS. Loadings were then summed across the SNP set to calculate the polygenic score.

The use of polygenic scores in population research is complicated by population stratification, or the non-random patterning of alleles across global populations (Cardon and Palmer 2003). Population stratification is a potential confound in genetic association studies (Hamer and Sirota 2000) and, by extension, polygenic score analysis (D. W. Belsky and Israel 2014). GWAS and polygenic scoring rely on a subset of genetic markers to act as proxies for unmeasured genetic variation. This approach is effective because much of the genome is in what is called “linkage disequilibrium,” i.e. certain groups genotypes tend to be inherited together and are thus correlated. Measuring one in a set of such genotypes can effectively provide information about the multiple unmeasured genotypes nearby. However, because patterns of linkage disequilibrium reflect genetic inheritance, they vary across populations of different ancestry. A result of ancestry-associated differences in patterns of linkage disequilibrium is that a given genotype may contain different information about the genome when measured in one population as compared to another (Carlson et al. 2013; Shifman et al. 2003). In the context of polygenic scoring, applying GWAS results derived in one population to compute a polygenic score in a different population introduces measurement error. This is one reason why polygenic scores derived from GWAS of European populations have reduced effect sizes in African American populations (D. W. Belsky et al. 2013; Domingue et al. 2014, 2015).

Importantly, reduction in the predictive accuracy of polygenic scores may also reflect confounding by environmental factors that are differentially distributed across populations. It is possible that large differences in environmental factors such as neighborhood social disorganization or exposure to discrimination suppress the influence of genetic factors in minority populations compared to non-Hispanic and white populations (Boardman et al. 2012, 2017). We believe that these are critical questions; however, given differences in minor-allele frequency and linkage disequilibrium patterns across groups, the purpose of this study – testing gene-environment correlation – is best served by restricting the analytic sample to increase the comparability to the population studied in the original GWAS. We therefore restrict primary analysis to unrelated women self-reporting white and non-Hispanic race/ethnicity.

Sample restrictions based on self-reported race/ethnicity may not completely protect against population stratification-related confounding (C. D. Campbell et al. 2005). To address residual population stratification within self-reported non-Hispanic whites in Add Health, we follow established practice and adjust analyses for principal components estimated from the genome-wide SNP data (Price et al. 2010). We estimate principal components among non-Hispanic white respondents using the genome-wide SNP data according to the method described by Price and colleagues (2006) using the PLINK command *pca*. We then residualize polygenic scores for the first ten principal components, i.e. we regressed polygenic scores on the ten principal-component scores and computed residual values from the predictions (Conley, Laidley, Belsky, et al. 2016). Residualized polygenic scores are standardized (mean = 0, standard deviation = 1) for analysis.

We scaled the polygenic score for age at menarche so that higher values correspond to genetic prediction of earlier age at menarche. We hereafter refer to this measure as the *accelerated menarche polygenic score*. Similarly, we scaled the polygenic score for age at first birth so that higher values correspond to genetic prediction of earlier age at first birth. We hereafter refer to this measure as the *earlier first-birth polygenic score*. Accelerated menarche and earlier first-birth polygenic scores are computed for n=2,631 women.

### Methods

We use survival methods to analyze timing of reproductive events. We employ the Kaplan-Meier method of estimating the survival function to describe the cumulative proportion of females experiencing an event by a given age. This method accommodates censoring, where a woman has not yet experienced an event by the end of follow-up, and provides estimates of mean age at event as the area under the curve. We use the Mantel-Haenszel log-rank test to compare survival curves for different risk groups as a way to test for the association of covariates.

We use a non-parametric proportional hazard model approach to test associations of genetic and environmental risks with accelerated reproductive development. This approach yields coefficients with a relative hazard interpretation, similar to the more familiar Cox model. The non-parametric proportional hazard approach can accommodate ties in event times, which are common in discretely measured data such as age-at-menarche or age-at-first-birth.

We estimate a generalized linear model with binomial error structure and a complementary log-log link function. The model is expressed as 
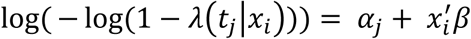
where *λ*(*t_j_* | *x_i_*) is the hazard of individual *i* at time *t_j_*, and *α_j_* is the complementary log-log transformation of the baseline hazard *λ_0_*(*t_j_*). This method accommodates continuous time measured in discrete intervals (completed years), and does not make any assumptions about the shape of the baseline hazard or the hazard in the interval. Individuals only contribute exposure intervals with complete information; for example, a respondent who reports 12 as age at menarche could have experienced the event any time between her 12^th^ and 13^th^ birthday. However, because we cannot know when exactly in the interval the event occurred, she contributes exposure only to the preceding intervals (i.e., up to age 12).

## Results

### Father Absence and Reproductive Timing

Table 2 presents hazard ratios for the relationship between father absence and reproductive development. Childhood experience of father absence – from birth to age seven – is significantly associated with earlier timing of all three reproductive events. At every age, the risk of menarche is 20% higher, on average, for girls who experienced father absence in childhood compared to peers who lived continuously with their fathers from birth to age seven. Exposure to father absence by age seven is also associated with earlier age at first sex and first birth, with larger effect sizes (hazard ratio (HR) = 1.42 and 1.48, respectively).

**Table 2.**
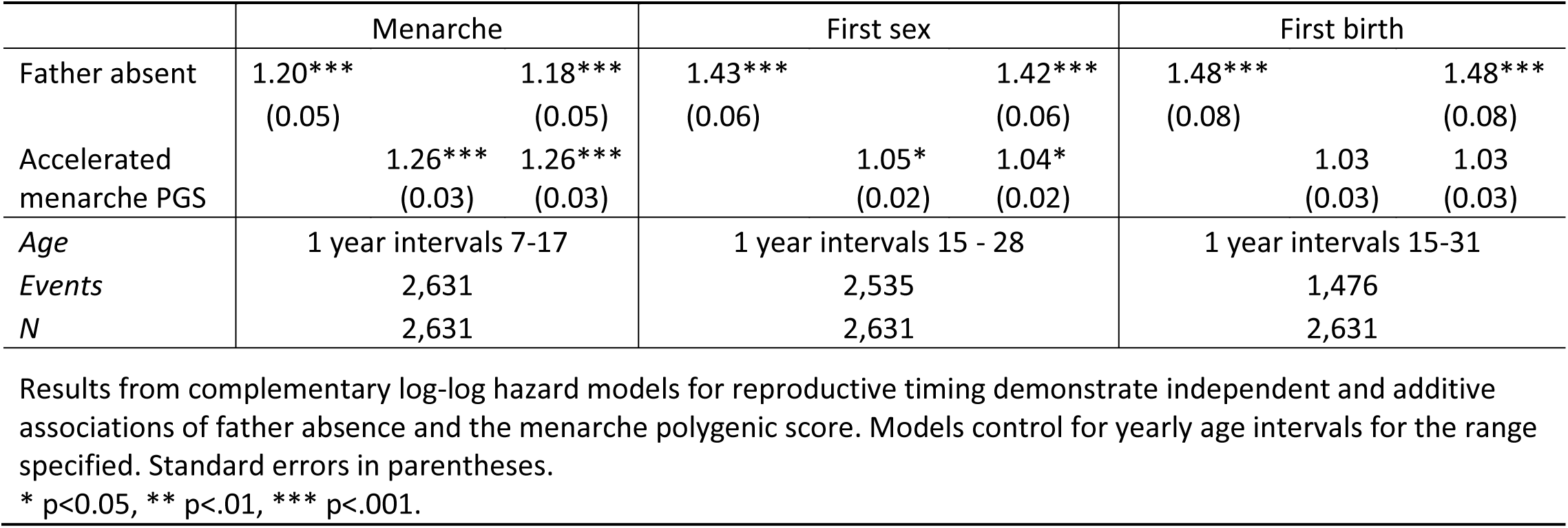
Hazard ratios for the risk of reproductive events and accelerated menarche polygenic score (PGS): Unrelated white females.

We illustrate the magnitude of the associations between father absence and reproductive timing in Figure 2. Across all reproductive events, father absence is associated with a significant acceleration in timing. Girls with absent fathers experience menarche earlier than girls with present fathers (12.1 versus 12.3, respectively). Similarly, girls with absent fathers experience sexual debut almost a full year earlier compared to those with present fathers (16.3 versus 17.8, respectively). Finally, girls with absent fathers go on to have their first birth nearly two years earlier on average compared to those with present fathers (24.9 and 26.7, respectively).

**Figure 2.**
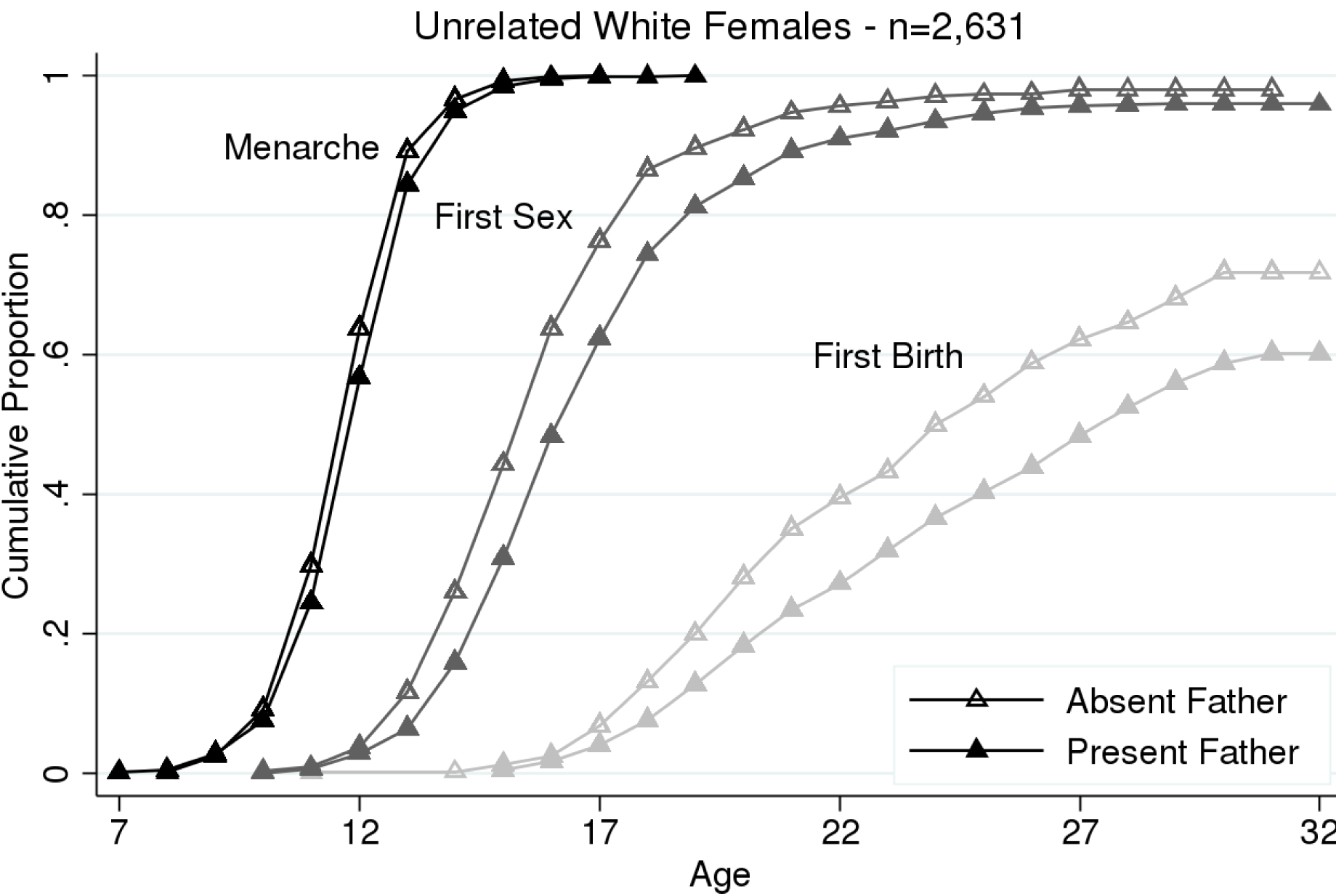
Reproductive Timing by Father Absence. Mantel-Haenszel log-rank test indicates that the differences between reproductive timing for females with absent and present fathers are statistically significant for all reproductive events (p<.001). Mean time to event: menarche, absent = 12.1, present = 12.3; first sex, absent = 16.3, present = 17.6; first birth, absent = 24.9, present = 26.7

### Accelerated Menarche Polygenic Score

Girls with higher accelerated menarche polygenic scores experience menarche at younger ages, on average, compared to peers with lower polygenic scores (HR = 1.26, Table 2). Similarly, women wither accelerated menarche polygenic scores experience sexual debut earlier (HR = 1.05). There is no association between the accelerated menarche polygenic score and age at first birth.

Figure 3 presents the cumulative proportion of women experiencing each reproductive event by age separately for high and low polygenic score groups. High score (solid) is defined as greater than or equal to one standard deviation above the mean, and low score (hollow) is defined as less than or equal to one standard deviation below the mean. Girls in the high polygenic score group experience menarche almost a full year earlier on average, compared to those in the low polygenic score group (11.8 versus 12.7, respectively). Similarly, those in the high polygenic score group also have earlier sexual debut, approximately four months ahead of those in the low polygenic score group (17.1 versus 17.4, respectively).

**Figure 3.**
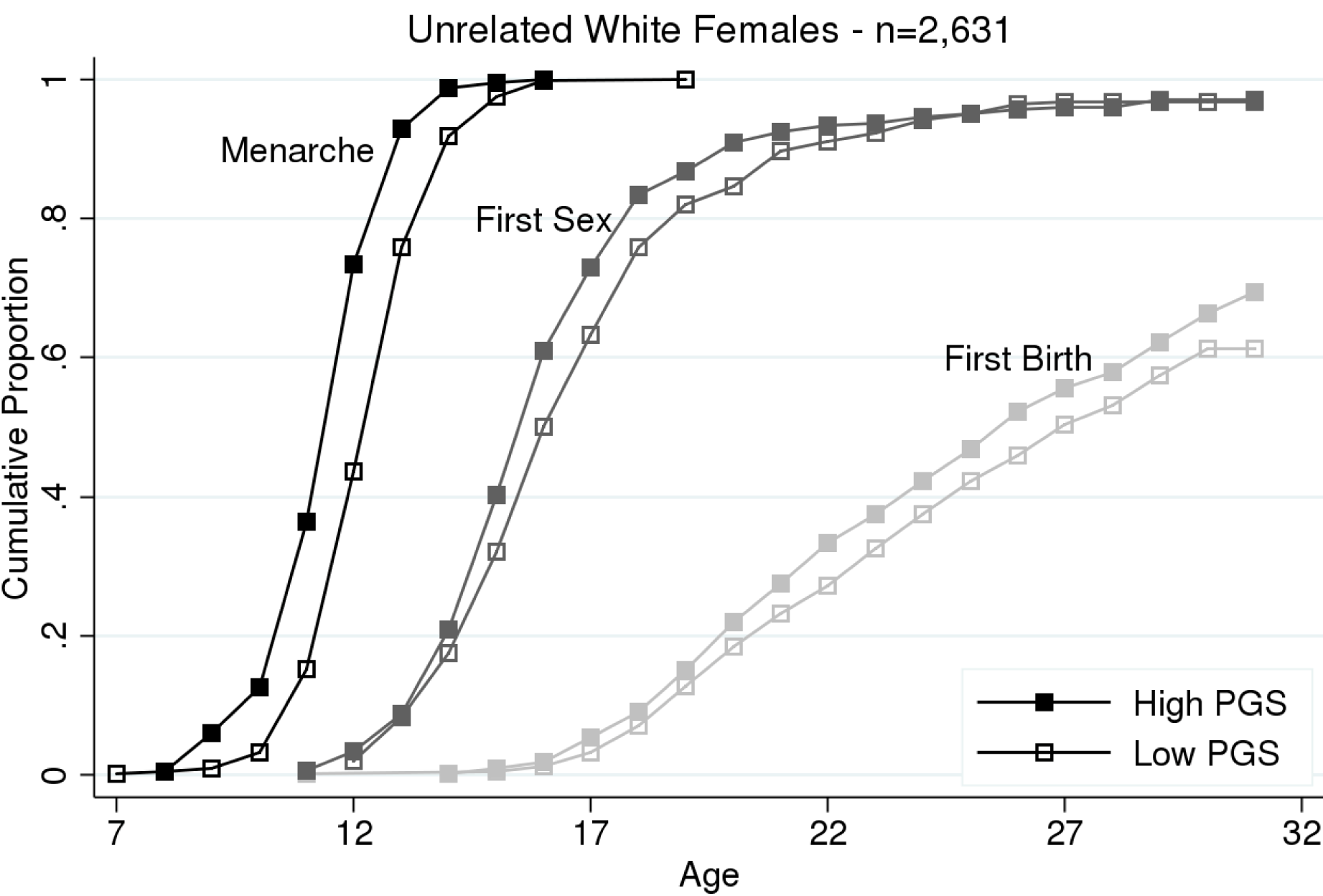
Reproductive Timing by Accelerated Menarche PGS. High PGS ≥ 1SD above mean (n=404); low PGS ≤ 1SD below mean (n=409). Mantel-Haenszel log-rank test indicates that the difference between high and low PGS groups is statistically significant for menarche (p<.001) and first sex (p<.05). Mean time to event: menarche, high = 11.8, low = 12.7; first sex, high = 17.1, low = 17.4; first birth, high = 25.8, low = 25.9

### Testing Gene-Environment Correlation

We test for gene-environment correlation in the relationship between father absence and menarcheal timing by computing correlations between girls’ accelerated menarche polygenic scores and exposure to father absence. The accelerated menarche polygenic score is not associated with father absence; there is no correlation between the accelerated menarche polygenic score and father absence (r=.01, Table 1). In Table 3, we compare the mean standardized polygenic scores by father absence; girls with absent fathers have similar accelerated menarche polygenic scores to those with present fathers.

**Table 3.**
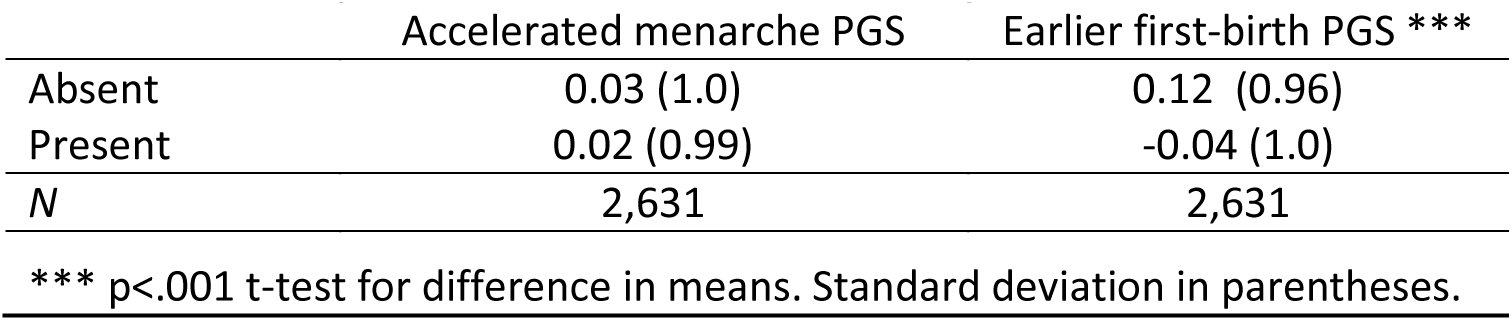
Mean polygenic score (PGS) by father coresidence: Unrelated white females.

|  | Accelerated menarche PGS | Earlier first-birth PGS *** |
| --- | --- | --- |
| Absent | 0.03 (1.0) | 0.12 (0.96) |
| Present | 0.02 (0.99) | -0.04 (1.0) |
| *N* | 2,631 | 2,631 |

\*\*\* $p < .001$ t-test for difference in means. Standard deviation in parentheses.

We formally test for confounding by gene-environment correlation in a hazard model regressing father absence on reproductive timing, controlling for the accelerated menarche polygenic score (Table 2). Father absence remains significantly associated with age at menarche, first sex, and first birth after controlling for known genetic influence on menarcheal timing (Table 2). The magnitude of these associations are unchanged. For age at menarche in particular, father absence and genetic risk are independent and additive predictors. There is no evidence that the genetics of menarche confound the relationship between father absence and pubertal timing.

### Earlier First-Birth Polygenic Score

We conduct the same analyses as outlined above using the polygenic score for age at first birth. On average, women with higher earlier first-birth polygenic scores experience accelerated timing of all three reproductive events (Table 4). Similar to results for the accelerated menarche polygenic score, the effect size for the standardized earlier first-birth polygenic score is largest for the phenotype matching the original GWAS (for age at menarche, HR = 1.08; for age at first sex, HR=1.21; for age at first birth, HR=1.30).

**Table 4.**
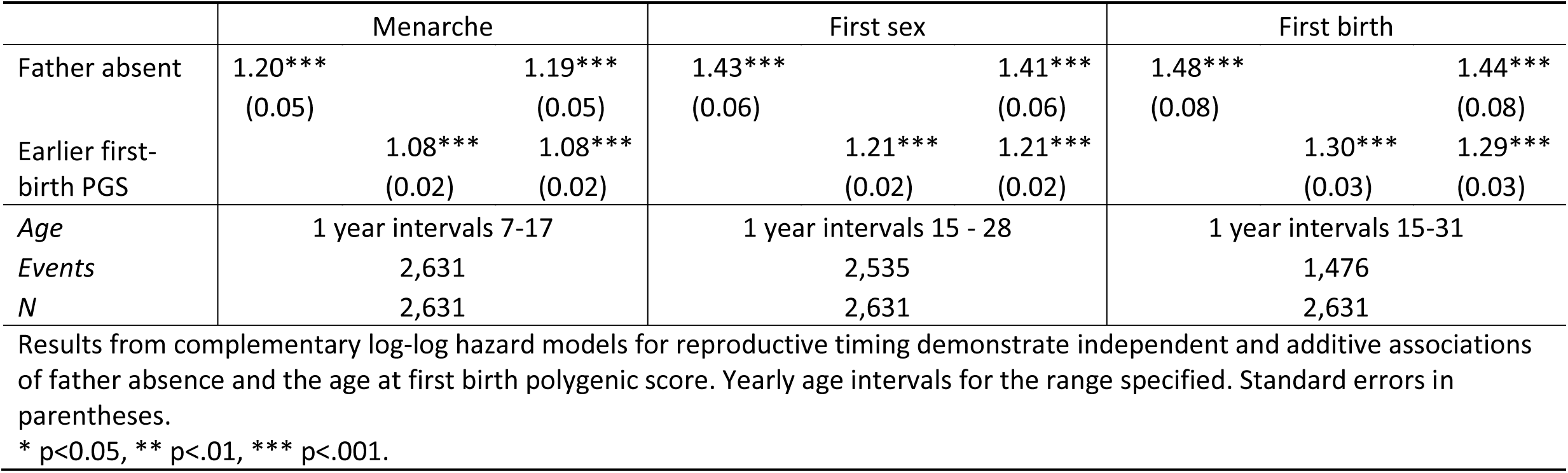
Hazard ratios for the risk of reproductive events and earlier first-birth polygenic score (PGS): Unrelated white females.

Figure 4 presents the cumulative proportion of women experiencing each reproductive event by age separately for high and low earlier first-birth polygenic score groups. High score (solid) is defined as greater than or equal to one standard deviation above the mean, and low score (hollow) is defined as less than or equal to one standard deviation below the mean. Girls in the high earlier first-birth polygenic score group experience menarche approximately four months earlier than their peers in the low polygenic score group (12.1 versus 12.4, respectively). The gap in reproductive timing widens for subsequent events; girls in the high earlier first-birth polygenic score group experience first sex over two years earlier, and first birth more than three years earlier, compared to girls in the low earlier first-birth polygenic score group.

**Figure 4.**
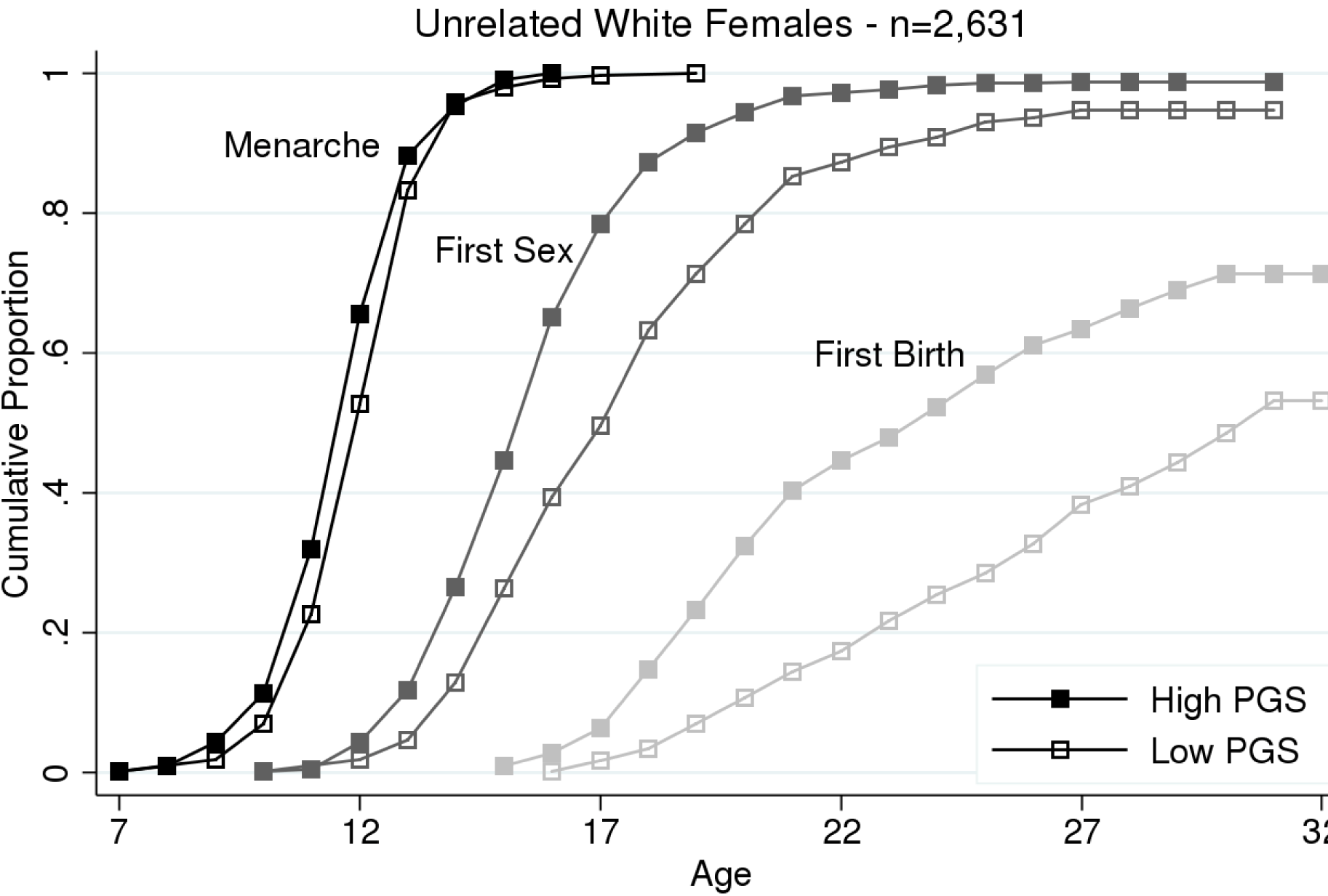
Reproductive Timing by Earlier AFB PGS. High PGS ≥ 1SD above mean (n=434); low PGS ≤ 1SD below mean (n=408). Mantel-Haenszel log-rank test indicates that the difference between high and low PGS groups is statistically significant for menarche (p<.01), first sex (p<.001), and first birth (p<001). Mean time to event: menarche, high = 12.1, low = 12.4; first sex, high = 16.2, low = 18.5; first birth, high = 24.7, low = 28.1

Finally, we test the gene-environment correlation hypothesis using the earlier first-birth polygenic score. The earlier first-birth polygenic score and father absence are correlated at r=.08 (Table 1); women who experience father absence have significantly higher earlier first-birth polygenic scores (Table 3). There is some evidence that the distribution of earlier first-birth polygenic scores varies by father absence, as shown in the right panel of Figure 5. However, we test whether the association between father absence and reproductive timing is confounded by the earlier first-birth polygenic score (Table 4). When the earlier first-birth polygenic score is added to the hazard models, effect-sizes for associations between father absence and reproductive timing phenotypes are slightly reduced, but remain statistically significant.

**Figure 5.**
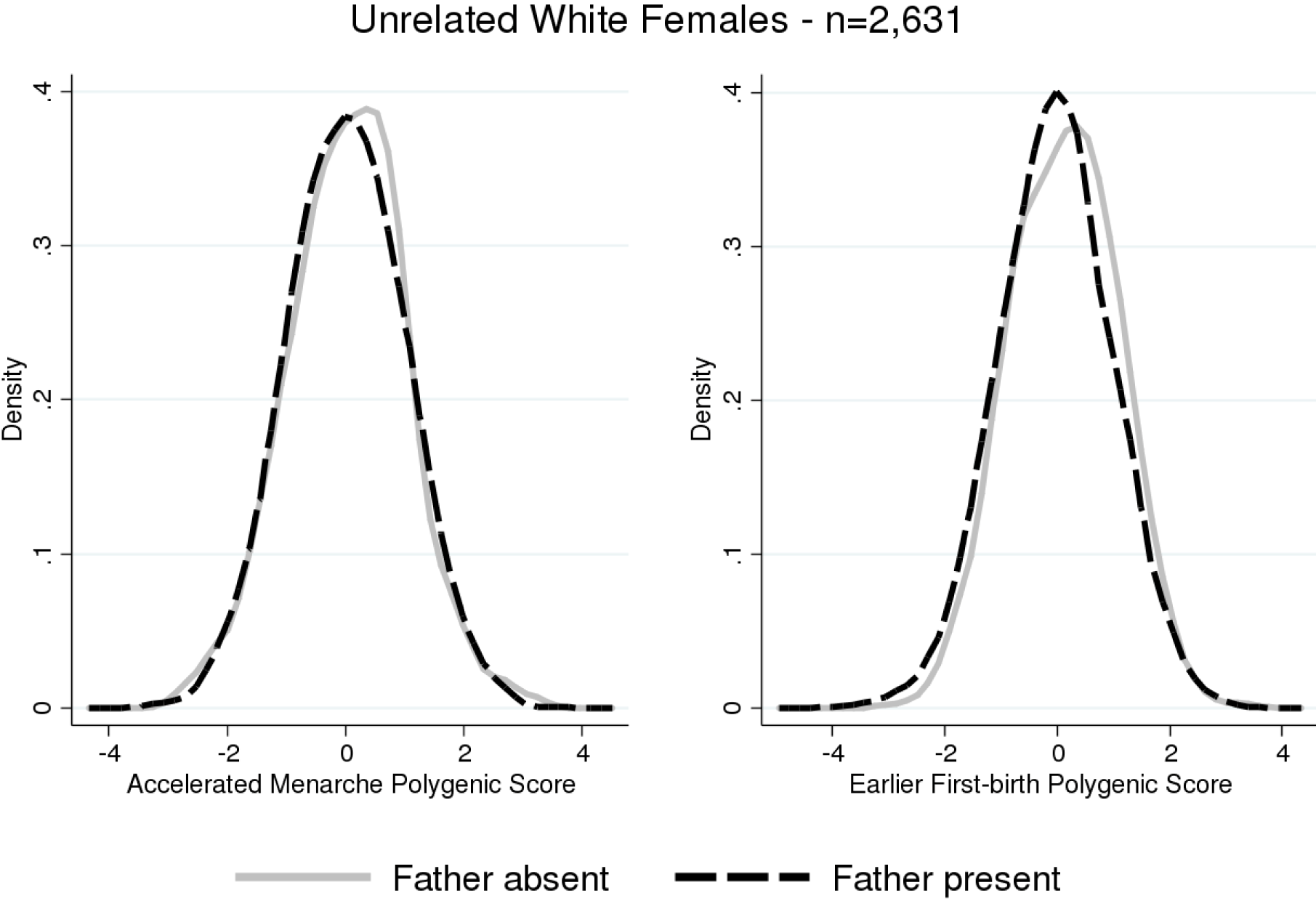
PGS Distribution by Father Absence. The distribution of polygenic scores by father absence tests for gene-environment correlation. The distribution of the accelerated menarche polygenic score is similar for girls in both groups. The distribution of the earlier first-birth polygenic score is shifted slightly to the right for girls who experience father absence.

### Robustness

As described in the methods section, our analysis focuses on non-Hispanic white females because the original GWAS were conducted on individuals of European ancestry. Consequently, the polygenic score constructed from the GWAS is expected to be less predictive among other ancestry groups. In Table 5, we present results from hazard models for non-Hispanic black and Hispanic females. As expected, the scores are less predictive in non-white samples, although there are modest associations across groups for the scores and their GWAS phenotypes.

**Table 5.** Hazard ratios for the risk of reproductive events and polygenic scores (PGS) by self-reported race/ethnicity.

|  | <u>Menarche</u> |  |  | <u>First sex</u> |  |  | <u>First birth</u> |  |  |
| --- | --- | --- | --- | --- | --- | --- | --- | --- | --- |
|  | White | Black | Hispanic | White | Black | Hispanic | White | Black | Hispanic |
| Accelerated menarche PGS | 1.26***<br>(0.03) | 1.09*<br>(0.04) | 1.32***<br>(0.06) | 1.05*<br>(0.02) | 0.99<br>(0.04) | 1.03<br>(0.04) | 1.03<br>(0.03) | 1.03<br>(0.05) | 1.03<br>(0.05) |
| Earlier first-birth PGS | 1.08***<br>(0.02) | 1.03<br>(0.04) | 1.12**<br>(0.05) | 1.21***<br>(0.02) | 1.08*<br>(0.04) | 1.10*<br>(0.04) | 1.30***<br>(0.03) | 1.14**<br>(0.05) | 1.12*<br>(0.06) |
| N | 2,631 | 796 | 656 | 2,631 | 796 | 656 | 2,631 | 796 | 656 |

Results from separate complementary log-log hazard models for reproductive timing for each polygenic score by self-reported race/ethnicity. Associations are in the same direction in the non-Hispanic black and Hispanic subsamples as in the non-Hispanic white analytic sample. Standard errors in parentheses. Yearly age intervals also controlled for in each model.
\* p<0.05, \*\* p<.01, \*\*\* p<.001.

Ultimately, the only way to exclude confounding to genetic analysis from population stratification is to analyze genetic differences between relatives – who share the same ancestry (Conley et al. 2015). We conducted our primary analysis using a subsample of unrelated individuals. Add Health also includes a sibling sample; we used this sample to repeat polygenic score analyses within a design that excludes potential confounding by population stratification. We tested genetic associations using sibling fixed effects models. We report results in Table 6. Among sisters, the carrier of the higher accelerated menarche polygenic score experienced earlier menarche as compared to her lower-scored sister. The effect-size was approximately the same as was observed in analysis of unrelated individuals, arguing strongly against confounding by population stratification. In contrast, the effect size for the earlier first-birth polygenic score was substantially reduced in the sibling fixed effects analysis and was no longer statistically significant. This reduction in effect size could signal some confounding by population stratification. Alternatively, it could imply other processes causing sisters to have correlated reproductive development outcomes. However, the sample size for sisters in the analysis of first birth is greatly reduced, as many women have not experienced a first birth by the time of last interview. Low discordance on father absence prohibits an analysis of father absence in the sister subsample.

**Table 6.** Comparing estimates from unrelated white female sample to sister sample.

| *Model* | Age at menarche |  |  |  | Age at first sex |  |  |  | Age at first birth |  |  |  |
| --- | --- | --- | --- | --- | --- | --- | --- | --- | --- | --- | --- | --- |
|  | OLS | FE | OLS | FE | OLS | FE | OLS | FE | OLS | FE | OLS | FE |
| Accelerated menarche PGS | -.32***<br>(0.03) | -0.27**<br>(0.09) |  |  | -.15*<br>(0.07) | -0.16<br>(0.17) |  |  | -1.12<br>(1.13) | -0.84<br>(4.01) |  |  |
| Earlier first-birth PGS |  |  | -.12***<br>(0.03) | -0.06<br>(0.09) |  |  | -.63***<br>(0.07) | -0.22<br>(0.17) |  |  | 7.46***<br>(1.15) | -6.08<br>(4.10) |
| *N* | 2,631 | 651 | 2,631 | 651 | 2,535 | 625 | 2,535 | 625 | 1,476 | 393 | 1,476 | 393 |

Results from separate linear regression models for reproductive timing for each polygenic score and reproductive event separately for unrelated white females and sister fixed-effects. Standard errors in parentheses.
\* $p < 0.05$ , \*\* $p < 0.01$ , \*\*\* $p < 0.001$ .

Results are robust to different measures of family environment (Table 7). Restricting the measure of father absence to non-residence at the time of birth, results are almost identical. Step-father coresidence is similarly predictive of earlier reproductive timing, and is not confounded by genetic risk.

**Table 7.**
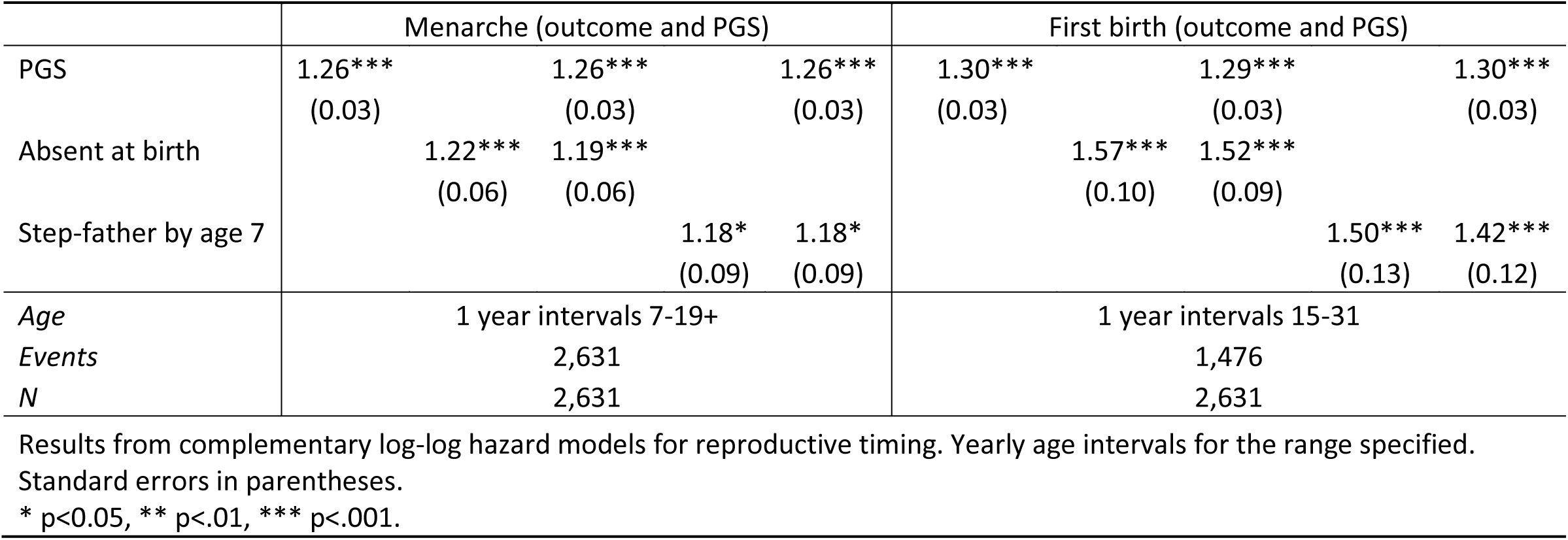
Alternative environmental exposures - hazard ratios for reproductive events: Unrelated white females.

### Limitations

This study has limitations. Ages at reproductive events are measured in years, which is imprecise, especially for timing of menarche. Data on age at first birth are right-censored, with follow-up ending between ages 24-32 (Add Health Wave IV). Although the modeling strategy accommodates this type of data, future analyses should consider women with completed reproduction. Father coresidence is reported retrospectively. However, a strength of our measure is that it draws information from both primary caregiver and child reports. Polygenic scores based on current GWAS results are incomplete measures of genetic influences on reproductive development. The GWAS we use to develop polygenic scores included hundreds of thousands of individuals. Nevertheless, the majority of genetic influence on reproductive development inferred based on family studies remains unexplained. As GWAS sample sizes increase, polygenic score performance is expected to improve (Okbay et al. 2016), which may change our ability to detect genetic confounding of associations between father absence and reproductive timing.

Residual population stratification may confound our analysis of gene-environment correlation. Our sibling comparison analysis rules out confounding by population stratification in the case of genetic associations with reproductive phenotypes. However, sibling comparison analysis was not feasible for tests of genetic association with father absence. We included statistical adjustment for principal components estimated from the genetic data to address this limitation, the standard in the field (Price et al. 2006). Nevertheless, studies with larger samples of siblings could strengthen confidence that our null findings for rGE were not confounded by residual population stratification.

Add Health represents a single cohort in the United States. Genetic influences on human traits and behaviors change over time and across space (Conley, Laidley, Boardman, et al. 2016; Demerath et al. 2013; Liu and Guo 2015; Tropf et al. 2016; Walter et al. 2016). Replication of our findings in samples from different birth cohorts and from different countries will clarify the extent to which findings from recent US birth cohorts generalize.

Finally, our findings are restricted to non-Hispanic white women. As we discuss above, this is motivated by the limitations of the current GWAS results, which are conducted in European ancestry populations. This is a pressing limitation to current empirical investigations of genetic and social factors influencing biological and behavioral outcomes, and GWAS in non-European ancestry populations should be a priority. It is our hope that new data, methods, or integrative frameworks become available in the near future that better enable researchers to utilize genome-wide data from all individuals.

## Discussion

We find no evidence of genetic confounding in the relationship between father absence and accelerated menarcheal timing. Women with higher accelerated menarche polygenic scores experience menarche and first sex earlier. Family structure risks for early puberty and sexual debut are independent of accelerated menarche polygenic scores. The rGE hypothesis explaining the link between father absence and earlier puberty is not supported in our sample. Father absence and GWAS-discovered genetic risk for pubertal timing are independent and additive predictors of adolescent development. If common genetic factors do link family environments and pubertal timing, they have not yet been uncovered in GWAS of menarche.

The findings for age at first birth are similar, although less conclusive. There is some evidence that girls with absent fathers have, on average, greater GWAS-discovered genetic risk for younger age at first birth compared to girls with present fathers. Although the magnitude of the associations between father absence and reproductive timing decline after controlling for the earlier first-birth polygenic score, the father absence association remains significant. Known genetic factors associated with age at first birth cannot account for the acceleration in reproductive timing associated with father absence.

Our findings do not support the use of molecular genetic testing in order to make specific predictions about outcomes for individuals. However, molecular genetic data can provide useful information about population parameters, contributing to our understanding of variation in reproductive timing. Polygenic scores for reproductive timing capture otherwise unobserved heterogeneity, allowing for more precise estimation of environmental effects, as well as investigations of interactions between genes and environments. We provide here an example of an approach proposed by Manski (2011) and others (D. W. Belsky and Israel 2014; Benjamin et al. 2012) that applies molecular genetic discoveries to test effects of environmental variables. Future studies may use the polygenic scores we studied as a measure of genetic liability to early puberty.

The lack of evidence of the genetic confounding hypothesis suggests that there may be something else underlying the consistent observation that girls who live apart from their fathers mature earlier than those who live together. While we are unable to test competing explanations, future research should investigate the possibility of psychosocial acceleration and the mechanisms through which biological embedding may occur, controlling for genetic confounds. Such research may identify targets for intervention to modify the reproductive trajectories on which family environments set developing girls.

## Acknowledgements

This research was supported by NRSA Individual Postdoctoral Fellowship NICHD 1F32HD084117-01, NICHD P2C-HD050924. This research uses data from Add Health, a program project directed by Kathleen Mullan Harris and designed by J. Richard Udry, Peter S. Bearman, and Kathleen Mullan Harris at UNC-CH, and funded by grant P01-HD31921 from the NICHD, with cooperative funding from 23 other federal agencies and foundations. DWB is an Early Career Fellow of the Jacobs Foundation and is supported by National Institute on Aging grants R21AG054846, R01AG032282, andP30AG028716.

## References

Alvergne, A., Faurie, C., & Raymond, M. (2008). Developmental plasticity of human reproductive development: Effects of early family environment in modern-day France. Physiology & Behavior, 95(5), 625–632. doi:10.1016/j.physbeh.2008.09.005

Anderson, K. G. (2015). Father Absence, Childhood Stress, and Reproductive Maturation in South Africa. Human Nature, 26(4), 401–425. doi:10.1007/s12110-015-9243-6

Barban, N., Jansen, R., de Vlaming, R., Vaez, A., Mandemakers, J. J., Tropf, F. C., et al (2016). Genome-wide analysis identifies 12 loci influencing human reproductive behavior. Nature Genetics, 48(12), 1462–1472. doi:10.1038/ng.3698

Belsky, D. W., & Israel, S. (2014). Integrating Genetics and Social Science: Genetic Risk Scores. Biodemography and Social Biology, 60(2), 137–155. doi:10.1080/19485565.2014.946591

Belsky, D. W., Moffitt, T. E., Sugden, K., Williams, B., Houts, R., McCarthy, J., & Caspi, A. (2013). Development and evaluation of a genetic risk score for obesity. Biodemography and social biology, 59(1), 85–100. doi:10.1080/19485565.2013.774628

Belsky, J., Steinberg, L. D., & Draper, P. (1991). Childhood experience, interpersonal development, and reproductive strategy: and evolutionary theory of socialization. Child Development, 62(4), 647–670. doi:10.1111/1467-8624.ep9109162242

Benjamin, D. J., Cesarini, D., Chabris, C. F., Glaeser, E. L., Laibson, D. I., Gu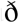nason, V., et al (2012). The Promises and Pitfalls of Genoeconomics. Annual Review of Economics, 4(1), 627–662. doi:10.1146/annurev-economics-080511-110939

Boardman, J. D., Barnes, L., Wilson, R., & Evans, D. (2012). Social disorder, APOE-E4 genotype, and change in cognitive function among older adults living in Chicago. Social Science & Medicine, 74(10), 1584–1590.

Boardman, J. D., McQueen, M. B., Wedow, R., Goode, J. A., Domingue, B. W., & Harris, K. M. (2017). Race differences in genetic associations for health and economic outcomes: the role of individual and institutional discrimination across the life course. Under Review.

Bogaert, A. F. (2008). Menarche and Father Avsence in a National Probability Sample. Journal of Biosocial Science, 40(04), 623–636. doi:10.1017/S0021932007002386

Booth, A., & Edwards, J. (1985). Age at marriage and marital instability. Journal of Marriage and the Family, 47(1), 67–75.

Browning, C., Leventhal, T., & Brooks-Gunn, J. (2004). Neighborhood context and racial differences in early adolescent sexual activity. Demography, 41(4), 697–720.

Bumpass, L. L., & Sweet, J. (1972). Differentials in marital instability: 1970. American Sociological Review, 37(6), 754–766.

Burt, S. A., McGue, M., DeMarte, J. A., Krueger, R. F., Iacono, W. G., H, P., et al (2006). Timing of Menarche and the Origins of Conduct Disorder. Archives of General Psychiatry, 63(8), 890. doi:10.1001/archpsyc.63.8.890

Bush, W. S., Moore, J. H., Li, J., McDonnell, S., & Rabe, K. (2012). Chapter 11: Genome-Wide Association Studies. PLoS Computational Biology, 8(12), e1002822. doi:10.1371/journal.pcbi.1002822

Campbell, B., & Udry, J. (1995). Stress and age at menarche of mothers and daughters. Journal of Biosocial Science, 27(2), 127–134.

Campbell, C. D., Ogburn, E. L., Lunetta, K. L., Lyon, H. N., Freedman, M. L., Groop, L. C., et al (2005). Demonstrating stratification in a European American population. Nature Genetics, 37(8), 868–872. doi:10.1038/ng1607

Cardon, L., & Palmer, L. (2003). Population stratification and spurious allelic association. The Lancet, 361(9357), 598–604.

Carlson, C. S., Matise, T. C., North, K. E., Haiman, C. A., Fesinmeyer, M. D., Buyske, S., et al (2013). Generalization and Dilution of Association Results from European GWAS in Populations of Non-European Ancestry: The PAGE Study. PLoS Biology, 11(9), e1001661. doi:10.1371/journal.pbio.1001661

Charalampopoulos, D., McLoughlin, A., Elks, C. E., & Ong, K. K. (2014). Age at Menarche and Risks of All-Cause and Cardiovascular Death: A Systematic Review and Meta-Analysis. American Journal of Epidemiology, 180(1), 29–40. doi:10.1093/aje/kwu113

Chisholm, J. S., Ellison, P. T., Evans, J., Lee, P. C., Lieberman, L. S., Pavlik, Z., et al (1993). Death, Hope, and Sex: Life-History Theory and the Development of Reproductive Strategies. Current Anthropology, 34(1), 1–24.

Comings, D. E., Muhleman, D., Johnson, J. P., & MacMurray, J. P. (2002). Parent-Daughter Transmission of the Androgen Receptor Gene as an Explanation of the Effect of Father Absence on Age of Menarche. Child Development, 73(4), 1046–1051. doi:10.1111/1467- 8624.00456

Conley, D., Domingue, B., Cesarini, D., Dawes, C., Rietveld, C., & Boardman, J. (2015). Is the Effect of Parental Education on Offspring Biased or Moderated by Genotype? Sociological Science, 2, 82–105. doi:10.15195/v2.a6

Conley, D., Laidley, T., Belsky, D. W., Fletcher, J. M., Boardman, J. D., & Domingue, B. W. (2016). Assortative mating and differential fertility by phenotype and genotype across the 20th century. Proceedings of the National Academy of Sciences of the United States of America, 113(24), 6647–52. doi:10.1073/pnas.1523592113

Conley, D., Laidley, T. M., Boardman, J. D., & Domingue, B. W. (2016). Changing Polygenic Penetrance on Phenotypes in the 20th Century Among Adults in the US Population. Scientific Reports, 6, 30348. doi:10.1038/srep30348

Culpin, I., Heron, J., Araya, R., Melotti, R., Lewis, G., & Joinson, C. (2014). Father absence and timing of menarche in adolescent girls from a UK cohort: The mediating role of maternal depression and major financial problems. Journal of adolescence, 37(3), 291–301.

Day, F. R., Perry, J. R. B. B., & Ong, K. K. (2015). Genetic regulation of puberty timing in Humans. Neuroendocrinology, 102(4), 1–13. doi:10.1159/000431023

Demerath, E. W., Choh, A. C., Johnson, W., Curran, J. E., Lee, M., Bellis, C., et al (2013). The positive association of obesity variants with adulthood adiposity strengthens over an 80- year period: a gene-by-birth year interaction. Human heredity, 75(2-4), 175–85. doi:10.1159/000351742

Domingue, B. W., Belsky, D. W., Conley, D., Harris, K. M., & Boardman, J. D. (2015). Polygenic Influence on Educational Attainment. AERA Open, 1(3), 233285841559997. doi:10.1177/2332858415599972

Domingue, B. W., Belsky, D. W., Harris, K. M., Smolen, A., McQueen, M. B., & Boardman, J. D. (2014). Polygenic Risk Predicts Obesity in Both White and Black Young Adults. PLoS ONE, 9(7), e101596. doi:10.1371/journal.pone.0101596

Draper, P., & Harpending, H. (1982). Father absence and reproductive strategy: An evolutionary perspective. Journal of anthropological research, 255–273.

Dudbridge, F., Visscher, P., Brown, M., McCarthy, M., Yang, J., Wray, N., et al (2013). Power and Predictive Accuracy of Polygenic Risk Scores. PLoS Genetics, 9(3), e1003348. doi:10.1371/journal.pgen.1003348

Elks, C. E., Perry, J. R. B., Sulem, P., Chasman, D. I., Franceschini, N., He, C., et al (2010). Thirty new loci for age at menarche identified by a meta-analysis of genome-wide association studies. Nature Genetics, 42(12), 1077–1085. doi:10.1038/ng.714

Ellis, B. J. (2004). Timing of pubertal maturation in girls: An integrated life history approach. Psychological Bulletin, 130(6), 920–958. doi:10.1037/0033-2909.130.6.920

Ellis, B. J., Bates, J. E., Dodge, K. A., Fergusson, D. M., Horwood, L. J., Pettit, G. S., & Woodward, L. (2003). Does father absence place daughters at special risk for early sexual activity and teenage pregnancy? Child Development, 74(3), 801–21.

Ellis, B. J., & Garber, J. (2000). Psychosocial antecedents of variation in girls’ pubertal timing: Maternal depression, stepfather presence, and marital and family stress. Child Development, 71(2), 485–501.

Ellis, B. J., McFadyen-Ketchum, S., & Dodge, K. (1999). Quality of early family relationships and individual differences in the timing of pubertal maturation in girls: a longitudinal test of an evolutionary model. Journal of personality and social psychology, 77(2), 387.

Ellis, B. J., Shirtcliff, E. A., Boyce, W. T., Deardorff, J., & Essex, M. J. (2011). Quality of early family relationships and the timing and tempo of puberty: effects depend on biological sensitivity to context. Development and psychopathology, 23(1), 85–99. doi:10.1017/S0954579410000660

Feng, Y., Hong, X., Wilker, E., Li, Z., Zhang, W., Jin, D., et al (2008). Effects of age at menarche, reproductive years, and menopause on metabolic risk factors for cardiovascular diseases. Atherosclerosis, 196(2), 590–597. doi:10.1016/j.atherosclerosis.2007.06.016

Foster, H., Hagan, J., & Brooks-Gunn, J. (2008). Growing up fast: Stress exposure and subjective “weathering” in emerging adulthood. Journal of health and social behavior, 49(2), 162–177. doi:10.1177/002214650804900204

Graber, J. A., Brooks-Gunn, J., & Warren, M. P. (1995). The Antecedents of Menarcheal Age: Heredity, Family Environment, and Stressful Life Events. Child Development, 66(2), 346. doi:10.2307/1131582

Hamer, D., & Sirota, L. (2000). Beware the chopsticks gene. Molecular Psychiatry, 5(1), 11–13. doi:10.1038/sj.mp.4000662

Hardy, J. B., Astone, N. M., Brooks-Gunn, J., Shapiro, S., & Miller, T. L. (1998). Like mother, like child: intergenerational patterns of age at first birth and associations with childhood and adolescent characteristics and adult outcomes in the second. Developmental psychology, 34(6), 1220.

Harris, K. M. (2010). An integrative approach to health. Demography, 47(1), 1–22.

Harris, K. M., Halpern, C. T., Hussey, J. M., Whitsel, E. a., Killeya-Jones, L. A., Tabor, J. W., et al (2013). Social, Behavioral, and Genetic Linkages from Adolescence Into Adulthood. American Journal of Public Health, 103(S1), S25–S32. doi:10.2105/AJPH.2012.301181

He, C., Kraft, P., Chen, C., Buring, J., Paré, G., Hankinson, S. E., et al (2009). Genome-wide association studies identify loci associated with age at menarche and age at natural menopause. Nature Genetics, 41, 724–728.

He, C., Zhang, C., Hunter, D. J., Hankinson, S. E., Buck Louis, G. M., Hediger, M. L., & Hu, F. B. (2010). Age at Menarche and Risk of Type 2 Diabetes: Results From 2 Large Prospective Cohort Studies. American Journal of Epidemiology, 171(3), 334–344. doi:10.1093/aje/kwp372

Highland, H., Avery, C., Duan, Q., Li, Y., & Harris, K. M. (2017). Quality Control Analysis of Add Health GWAS Data.

Hoier, S. (2003). Father absence and age at menarche. Human Nature, 14(3), 209–233. doi:10.1007/s12110-003-1004-2

Igra, V., & Irwin, C.Jr (1996). Theories of adolescent risk-taking behavior. In R. J. DiClemente, W. B. Hansen, & L. E. Ponton (Eds.), Handbook of adolescent health risk behavior (pp. 35–51). Springer US.

Karapanou, O., & Papadimitriou, A. (2010). Determinants of menarche. Reproductive Biology and Endocrinology, 8(115).

Kiernan, K. E. (1977). Age at puberty in relation to age at marriage and parenthood: A national longitudinal study. Annals of Human Biology, 4(4), 301–308. doi:10.1080/03014467700002241

Kiernan, K. E., & Hobcraft, J. (1997). Parental divorce during childhood: Age at first intercourse, partnership and parenthood. Population Studies, 51(1), 41–&. doi:10.1080/0032472031000149716

Lakshman, R., Forouhi, N. G., Sharp, S. J., Luben, R., Bingham, S. A., Khaw, K.-T., et al (2009). Early Age at Menarche Associated with Cardiovascular Disease and Mortality. The Journal of Clinical Endocrinology & Metabolism, 94(12), 4953–4960. doi:10.1210/jc.2009- 1789

Liu, H., & Guo, G. (2015). Lifetime Socioeconomic Status, Historical Context, and Genetic Inheritance in Shaping Body Mass in Middle and Late Adulthood. American Sociological Review, 80(4), 705–737. doi:10.1177/0003122415590627

Manski, C. F. (2011). Genes, eyeglasses, and social policy. The Journal of Economic Perspectives, 25(4), 83–93.

McQueen, M. B., Boardman, J. D., Domingue, B. W., Smolen, A., Tabor, J. W., Killeya-Jones, L. A., et al (2015). The National Longitudinal Study of Adolescent to Adult Health (Add Health) Sibling Pairs Genome-Wide Data. Behavior Genetics, 45(1), 12–23. doi:10.1007/s10519-014-9692-4

Mendle, J., Harden, K. P., Turkheimer, E., Van Hulle, C. a., D’Onofrio, B. M., Brooks-Gunn, J., et al (2009). Associations between father absence and age of first sexual intercourse. Child Development, 80(5), 1463–1480. doi:10.1111/j.1467-8624.2009.01345.x

Mendle, J., Ryan, R. M., & McKone, K. M. (2015). Early Childhood Maltreatment and Pubertal Development: Replication in a Population-Based Sample. Journal of Research on Adolescence, 26(3), 595–602. doi:10.1111/jora.12201

Mendle, J., Turkheimer, E., D’Onofrio, B. M., Lynch, S. K., Emery, R. E., Slutske, W. S., & Martin, N. G. (2006). Family structure and age at menarche: A children-of-twins approach. Developmental Psychology, 42(3), 533–542.

Moffitt, T., Caspi, A., Belsky, J., & Silva, P. (1992). Childhood experience and the onset of menarche: A test of a sociobiological model. Child Development, 63(1), 47–58.

Moore, S. R., Harden, K. P., & Mendle, J. (2014). Pubertal Timing and Adolescent Sexual Behavior in Girls. Developmental Psychology, 50(6), 1734–1745. doi:10.1037/a0036027

Okbay, A., Beauchamp, J., Fontana, M., Lee, J., & Pers, T. (2016). Genome-wide association study identifies 74 loci associated with educational attainment. Nature, 533(7604), 539–542.

Patton, G., & McMorris, B. (2004). Puberty and the onset of substance use and abuse. Pediatrics, 114(3), 300–306.

Perry, J., Day, F., Elks, C., Sulem, P., & Thompson, D. (2014). Parent-of-origin-specific allelic associations among 106 genomic loci for age at menarche. Nature, 514(7520), 92–97.

Plomin, R., DeFries, J., Knopik, V., & Neiderheiser, J. (2013). Behavioral genetics. Palgrave Macmillan.

Polderman, T. J. C., Benyamin, B., de Leeuw, C. a, Sullivan, P. F., van Bochoven, A., Visscher, P. M., & Posthuma, D. (2015). Meta-analysis of the heritability of human traits based on fifty years of twin studies. Nature Genetics, 47(7), 702–709. doi:10.1038/ng.3285

Price, A. L., Patterson, N. J., Plenge, R. M., Weinblatt, M. E., Shadick, N. A., & Reich, D. (2006). Principal components analysis corrects for stratification in genome-wide association studies. Nature Genetics, 38(8), 904–909. doi:10.1038/ng1847

Price, A. L., Zaitlen, N. A., Reich, D., & Patterson, N. (2010). New approaches to population stratification in genome-wide association studies. Nature Reviews Genetics, 11(7), 459–463. doi:10.1038/nrg2813

Quinlan, R. (2003). Father absence, parental care, and female reproductive development. Evolution and Human Behavior, 24(6), 376–390.

Remsberg, K. E., Demerath, E. W., Schubert, C. M., Chumlea, W. C., Sun, S. S., & Siervogel, R. M. (2005). Early Menarche and the Development of Cardiovascular Disease Risk Factors in Adolescent Girls: The Fels Longitudinal Study. The Journal of Clinical Endocrinology & Metabolism, 90(5), 2718–2724. doi:10.1210/jc.2004-1991

Rowe, D. C. (2000). Environmental and Genetic Influences on Pubertal Development: Evolutionary Life History Traits? In J. Rodgers, D. C. Rowe, & W. B. Miller (Eds.), Genetic Influences on Human Fertility and Sexuality (pp. 147–168). Kluwer Academic Publishers.

Rowe, D. C. (2002). On genetic variation in menarche and age at first sexual intercourse: A critique of the Belsky-Draper hypothesis. Evolution and Human Behavior, 23(5), 365–372. doi:10.1016/S1090-5138(02)00102-2

Ryan, R. M. (2015). Nonresident fatherhood and adolescent sexual behavior: A comparison of siblings approach. Developmental Psychology, 51(2), 211–223. doi:10.1037/a0038562

Sandler, D. P., Wilcox, A. J., & Horney, L. F. (1984). Age at menarche and subsequent reproductive events. American Journal of Epidemiology, 119(5), 765–774.

Shifman, S., Kuypers, J., Kokoris, M., Yakir, B., & Darvasi, A. (2003). Linkage disequilibrium patterns of the human genome across populations. Human Molecular Genetics, 12(7), 771–776. doi:10.1093/hmg/ddg088

Stoll, B. A., Vatten, L. J., & Kvinnsland, S. (1994). Does early physical maturity influence breast cancer risk? Acta oncologica, 33(2), 171–6.

Tamakoshi, K., Yatsuya, H., Tamakoshi, A., & Group, F. the J. S. (2011). Early age at menarche associated with increased all-cause mortality. European Journal of Epidemiology, 26(10), 771–778. doi:10.1007/s10654-011-9623-0

Tither, J. M., & Ellis, B. J. (2008). Impact of fathers on daughters’ age at menarche: A genetically and environmentally controlled sibling study. Developmental psychology, 44(5), 1409–1420.

Towne, B., Czerwinski, S. A., Demerath, E. W., Blangero, J., Roche, A. F., & Siervogel, R. M. (2005). Heritability of age at menarche in girls from the Fels Longitudinal Study. American Journal of Physical Anthropology, 128(1), 210–219. doi:10.1002/ajpa.20106

Trivers, R. (1972). Parental investment and sexual selection. In Sexual Selection & the Descent of Man (pp. 136–179). New York: Aldine de Gruyter.

Tropf, F. C., Verweij, R. M., Most, P. J. van der, Stulp, G., Bakshi, A., Briley, D. A., et al (2016). Mega-analysis of 31,396 individuals from 6 countries uncovers strong gene-environment interaction for human fertility. bioRxiv, 049163. doi:10.1101/049163

Udry, J. R. (2008). Age at menarche, at first intercourse, and at first pregnancy. Journal of Biosocial Science, 11(04), 433–441. doi:10.1017/S0021932000012517

Udry, J. R., & Cliquet, R. L. (1982). A Cross-Cultural Examination of the Relationship Between Ages at Menarche, Marriage, and First Birth. Demography, 19(1), 53. doi:10.2307/2061128

Walter, S., Mejía-Guevara, I., Estrada, K., Liu, S. Y., Glymour, M. M., & NA, C. (2016). Association of a Genetic Risk Score With Body Mass Index Across Different Birth Cohorts. JAMA, 316(1), 63. doi:10.1001/jama.2016.8729

Webster, G. D., Graber, J. A., Gesselman, A. N., Crosier, B. S., & Schember, T. O. (2014). A Life History Theory of Father Absence and Menarche: A Meta-Analysis. Evolutionary Psychology, 12(2), 147470491401200202. doi:10.1177/147470491401200202

Wu, L. L., & Martinson, B. C. (1993). Family structure and the risk of a premarital birth. American Sociological Review, 58(2), 210–232.

